# Efficient End-to-end Learning for Cell Segmentation with Machine Generated Incomplete Annotations

**DOI:** 10.1101/2022.07.03.498609

**Authors:** Prem Shrestha, Nicholas Kuang, Ji Yu

## Abstract

Automated cell segmentation from optical microscopy images is usually the first step in the pipeline of single-cell analysis. Recently, deep-learning based algorithms have shown superior performances for the cell segmentation tasks. However, a disadvantage of deep-learning is the requirement for a large amount of fully-annotated training data, which is costly to generate. Weakly-supervised and self-supervised learning is an active research area, but often the model accuracy is inversely correlated with the amount of annotation information provided. Here we focus on a specific subtype of incomplete annotations, which can be generated programmably from experimental data, thus allowing for more annotation information content without sacrificing the annotation speed. We designed a new model architecture for end-to-end training using such incomplete annotations. We benchmarked our method on a variety of publicly available dataset, covering both fluorescence and bright-field imaging modality. We additionally tested our method on a microscopy dataset generated by us, using machine generated annotations. The results demonstrated that our model trained under weak-supervision can achieve segmentation accuracy competitive to, and in some cases surpassing, state-of-the-art models trained under full supervision. Therefore, our method can be a practical alternative to the established full-supervision methods.

## Introduction

In the recent literature, deep-learning based cell segmentation methods have demonstrated unparalleled segmentation accuracy^1–6^ and are increasingly being adopted by biomedical researchers as the method of choice for microscopy image analysis^7–9^. However, a well-known disadvantage of the deep-learning methods is that they are very data-hungry, and require a large amount of annotated training data in order to achieve high model performance. For example, a recent study showed that the accuracy of the cell segmentation models did not reach saturation even after training with >1.6 million cell instances^10^. Training data for cell segmentation are particularly costly to generate, because the annotations have to be produced at the instance level, i.e, the exact boundaries of each cell need to be manually determined, unlike many other machine vision tasks such as image classification, where the annotations were produced at the image level. This drawback with regard to the training data further raises concerns over the scalability of the deep-learning method for three-dimensional microscopy, which is a highly active arena for modern biomedical research, but is even more difficult to produce manual annotations.

To alleviate the burden of image annotation, a significant amount of effort has been put into the studies of weakly-supervised learning^11^ and self-supervised learning^12^ from images. For example, within the general instance segmentation literature, bounding-box^11^, scribble^13^ and point^14,15^ annotations have been proposed as alternatives to the detailed instance masks. Advances have also been made in self-supervised learning^16–19^, which focuses on learning a useful representation of the image features without focusing on a specific task or using any labels; the learned representations can then be used for more specific downstream tasks, including image segmentation. For biomedical image segmentation, it has been proposed that class-activation maps^20^ can be utilized as a stand-in for segmentation masks^21–23^, because high gradients tend to localize to regions of importance for the modeling objective. This allows the model to be trained on simpler auxiliary tasks, such as image classification, which requires simpler annotation. In addition, for specific microscopy modalities, novel model architecture that intelligently incorporates the prior knowledge regarding the input image data is a successful strategy to reduce/eliminate the need for annotations^24^.

Faster annotation and higher model accuracy, however, are often two conflicting goals: the model accuracy generally improves with the more detailed annotations, which requires longer time to generate, and vice versa. Therefore, in this paper, we will focus on cell segmentation models trained with weak annotations that potentially can be produced programmably or semi-programmably from experimental data. We believe this strategy is the best compromise between the goals of high model accuracy and lesser annotation efforts (Fig. 1a). Specifically, we will focus on two types of annotations: (1) Image-level segmentations (Fig. 1b), which separates the cellular region from the background region. Such an annotation can be produced by acquiring a fluorescence image of the cellular sample and thresholding the resulting image. (2) Location-of-interests (LOIs), which are the rough locations for each cell present in the image (Fig. 1b). This annotation can be produced by acquiring nucleus images using widely employed labels, such as diamidino-2-phenylindole (DAPI). Nucleus detection from DAPI images is a heavily researched topic. Algorithms yielding accurate results are readily available. By focusing on these two specific annotations, a researcher can potentially generate a large dataset fairly quickly, and still offer annotations with rich informational content for training an accurate segmentation model.

**Figure 1.**
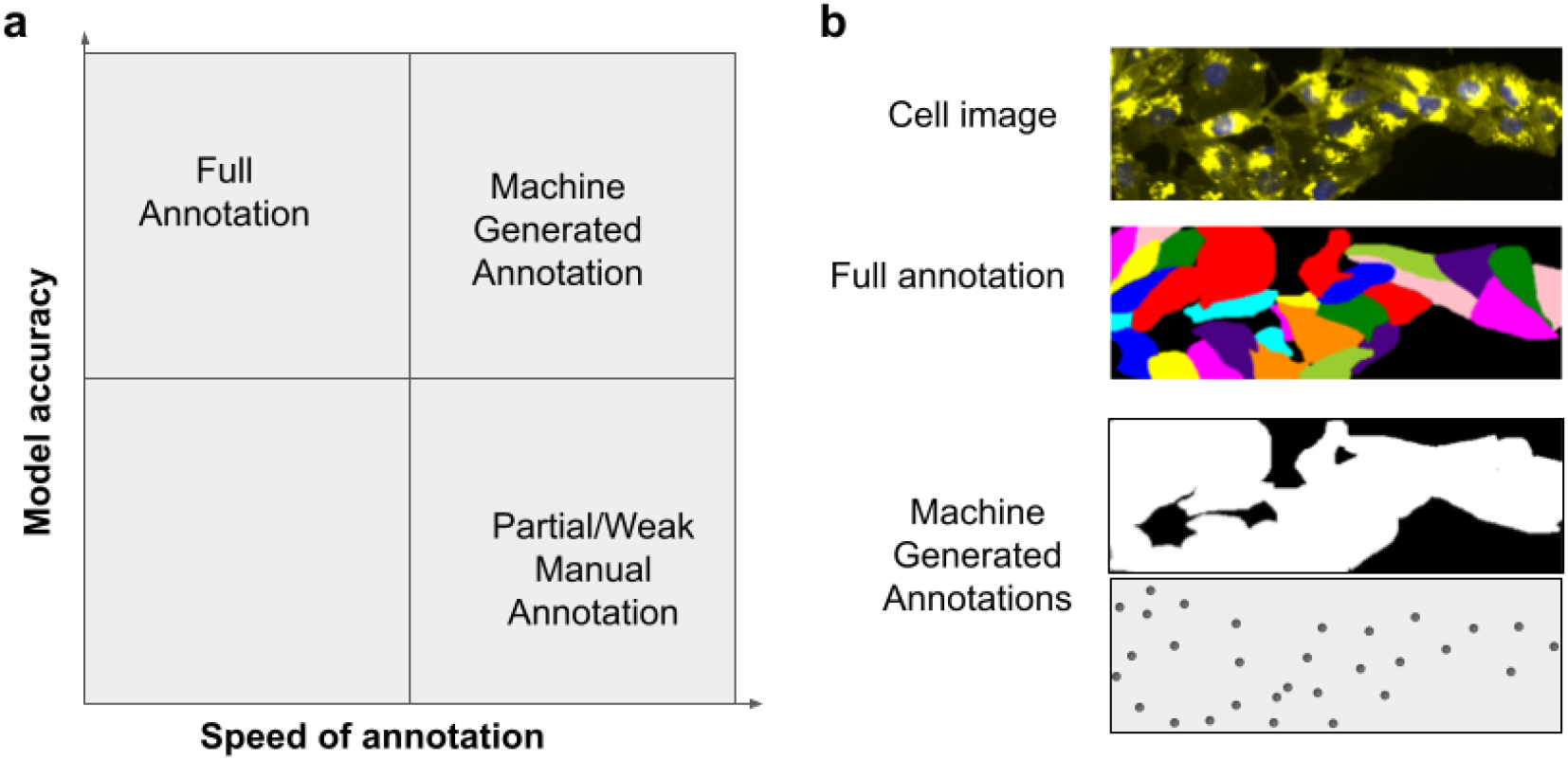
Comparison of various training data annotations for single-cell segmentation models. (**a**) Diagram illustrating the conflicting goals of faster annotation and higher model accuracy, and how experimentally acquired annotations offer a better compromise. (**b**) Examples of annotations. The top image shows a representative microscopy image of cells. The second row shows the corresponding full annotation of single-cell segmentations, which is expensive to generate. The bottom two rows showed two types of incomplete annotations that can potentially be generate programmably: (1) image-level segmentation, which labels the pixels of all cells in the image, and (2) location-of-interests annotation, which a subtype of the point annotation that denotes rough locations of individual cells.

## Results

### Model design

Fig. 2a shows our model architecture for the single-cell segmentation task. The overall model contains three sub-modules, all of which are based on fully-convolutional neural network (FCN) design. The backbone employs a standard encoder/decoder structure, and outputs feature representations extracted from the input image at multiple resolutions. The second component, the location proposal network (LPN), is analogous to the region proposal network (RPN) of the popular FastRCNN model for object detection^25^. Its task is to predict a set of LOIs from the feature outputs of the backbone. Unlike RPN, which computes regression for the object bounding-box (location and size), the LPN does not predict the object size, because the information is unavailable in the annotations. The last component, the segmentation FCN, is responsible for output the single-cell segmentations, taking the high-resolution image features as the input. To allow the network to focus on a specific cell, the image features were first concatenated with small tensors representing the predicted LOIs, one for each cell. This way, the segmentation FCN receives different inputs for different cells, and thus is able to produce variable segmentations even though the image features are constant. To increase the algorithm efficiency, the segmentation FCN only performs computation within a small area around the LOI, as it can reasonably assume that pixels far away from the LOI are not part of the segmentation (see supplementary text for implementation details). Combined, the three modules learn the segmentation task in an end-to-end manner. We name our network architecture location assisted cell segmentation system (LACSS).

**Figure 2.**
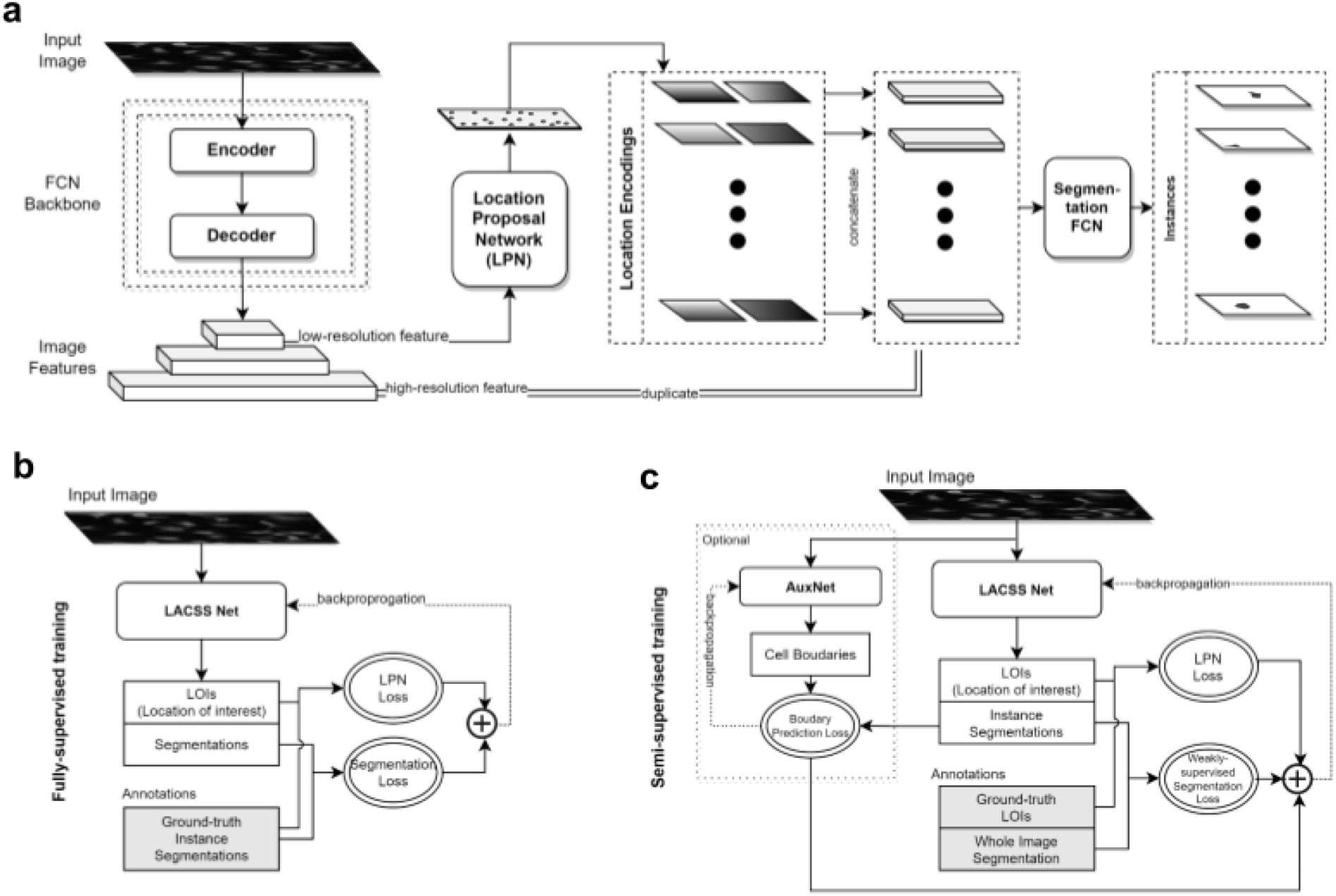
LACSS model. (**a**) Model architecture showing all three components of the model: backbone, location proposal network(LPN) and segmentation FCN. (**b**) Schematic diagram of LACSS model training under full-supervision. (**c**) Schematic diagram of LACSS model training under semi-supervision.

Even though the model is designed with incomplete annotation in mind, it is actually also able to learn under full supervision (Fig 2b). Under such a configuration, the model loss is the sum of two components. The first is the LPN loss, which measures the differences between the predicted LOIs and the ground truth LOIs. The latter can be computed from the individual cells’ ground truth segmentations. The second is the segmentation loss, which is simply the cross-entropy loss between the predicted cell segmentations and the ground truth cell segmentations. On the other hand, if the model is trained under the configuration of the weak supervision (Fig 2c), the LPN loss can still be calculated, because the LOIs are part of the image annotation. The segmentation loss, on the contrary, is no longer available. Therefore, we used instead a weakly-supervised segmentation loss function (see supplementary text), which computes (i) the consistency between single-cell segmentation with the image-level segmentation, and (ii) the consistency between individual cell segmentations. The latter part aims to minimize overlaps between individual cell segmentations. Detailed mathematical formulations of all loss functions are provided in supplementary text.

### Cell image library dataset

As a proof of principle, we first performed a quick segmentation test on the publicly available Cell Image Library^26^ dataset. The dataset consists of 100 images of dual channel (cytoplasm/nucleus) fluorescence cell images from a single experiment (Fig. 3). The images are of high signal-to-noise ratio and both cell-cell contacts and cell-cell occlusions are rare. Therefore segmentation is relatively easy. We split the data into training(n=89) and validation (n=11) sets, following Stringer et al.^2^| and performed model training using the weak-supervision configuration. The original dataset was pre-annotated with full segmentation masks. We converted the original cell segmentations to image-level segmentations by combining all cell masks, and used the center-of-mass of the individual cells as the LOIs.

**Figure 3.**
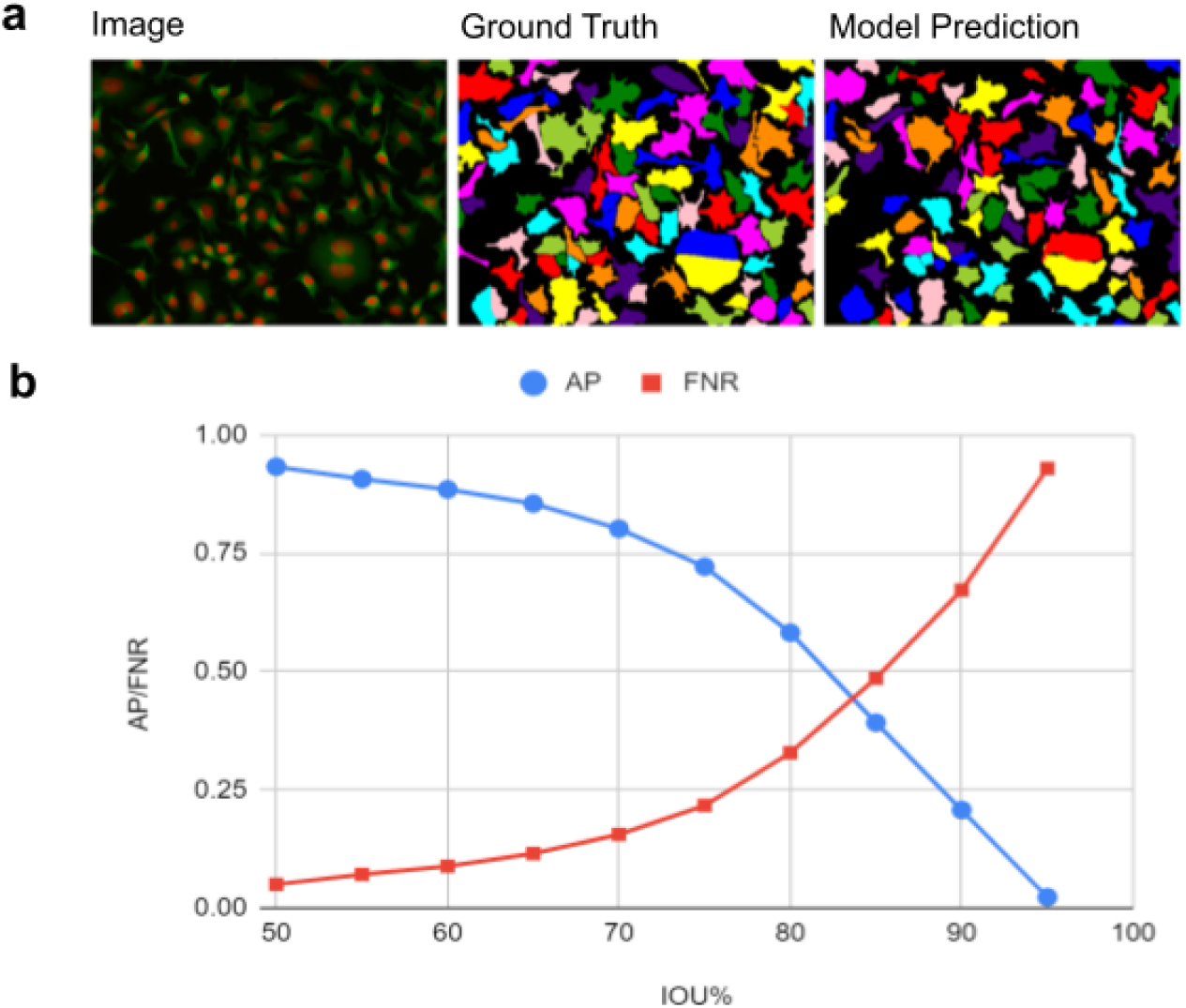
Segmentation of cell image library dataset with a LACSS model trained with incomplete annotations. (**a**) A representative example of segmentation results showing the input image (left), the ground-truth segmentation (middle) and model prediction (right). The pseudo colors were generated via skimage’s label2rgb function for visual clarity and carried no extra meaning. (b) Model performances quantified by average precisions and false negative rates at various IOU thresholds.

To benchmark the segmentation accuracy, we computed average precision (AP) on the validation set, as well as the false negative rate (FNR) as Edlund et.al^10^. Both metrics were computed under a series of intersection-over-unions (IOUs) criteria ranging from 50% to 95%. The model trainings were run with random initialization for multiple times (n=5). We found the best AP_50_ score surpassed 0.93 in all runs, with the best model producing AP_50_=0.933. The complete benchmarks of the best model is shown in Fig 3b.

Both the quantitative benchmarks and visual inspection of the segmentation results (Fig 3a) suggested that our model performs well on this dataset. To our surprise though, our results seem to be even better than previous reports based on segmentation models training with full supervision^2^. Using the exact same dataset, Stringer et al reported precisions (IOU=50%) from three different segmentation models (cellpose^2^, MaskRCNN^27^s and stardist^28^), all of which are lower than ours by more than ten percent points. We suspect this is simply due to insufficient optimization of the previous models. When we trained LACSS with full supervision, we obtained essentially the same accuracy (best AP_50_=0.937). However we did observe a strong tendency to overfit under full-supervision probably due to the small size of the dataset, which may have contributed to the poorer results reported previously.

### LIVECell dataset

Encouraged by the positive outcome on the cell image library data, we then moved on to test our model performances on a much larger dataset, LIVECell^10^. This is a recently published cell segmentation dataset containing >1.6 million fully segmented cells from eight different cell lines. The imaging modality is phase contrast, which has lower contrast than fluorescence for the purpose of cell detection. The cell lines chosen cover a diverse set of morphologies. Cell-cell contact and cell-cell occlusion are of frequent occurrences. Therefore, this is a more difficult study case for cell segmentation, but is also more representative of real-life problems in biomedical research. In addition, the dataset was pre-splitted into training, validation and testing sets, which allows for more useful benchmarking, and baseline benchmarks were already provided based on two state-of-the-art models, MaskRCNN^27^ and CenterMask^29^, trained on full-supervision.

Similar to the Cell Image Library case, we first converted the original cell-level annotations to image-level segmentations and LOIs. We then trained LACSS models with the converted weak annotations. We also train LACSS with full-supervision using the original annotation in order to understand the performance gap between the two configurations. We found that indeed LACSS trained under weak-supervision underperform by a quite large margin (Fig. 4). Among the eight cell lines, the SkBr3 line exhibited the smallest performance gap (AP_50_=0.857 / 0.948 for weakly-supervised/fully-supervised models), while the SHSY5Y line showed the largest gap (AP_50_=0.128 / 0.520).

**Figure 4.**
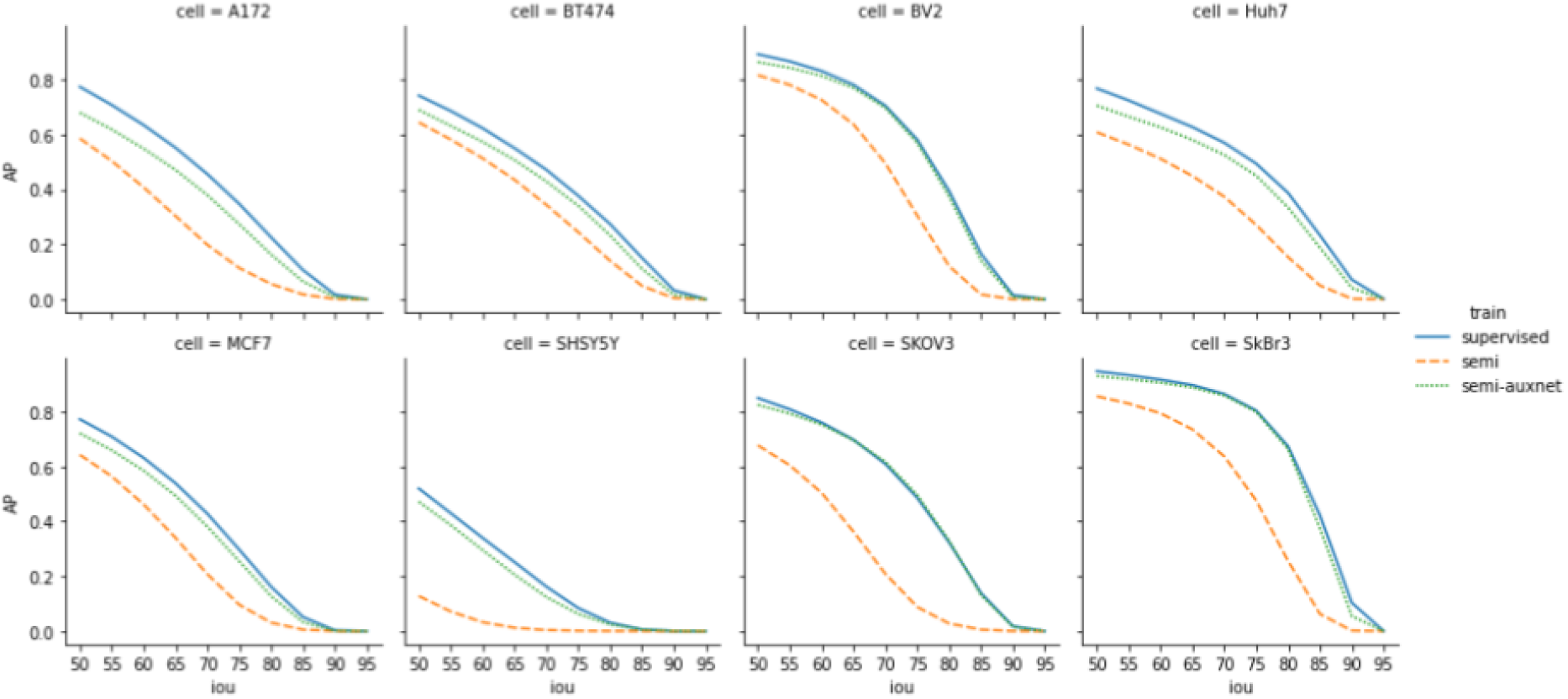
LACSS model accuracy trained with LIVECell dataset. Models were trained either under full-supervision using the original single-cell annotation (supervised, solid blue line), under semi-supervision using synthetic annotation but without auxnet (semi, dash line), or under semi-supervision with boundary-defining auxnet (semi-auxnet, dotted line). AP values were computed for eight cell lines at increasing IOU thresholds from 50% to 95%. The results showed significant improvements in model accuracy by introducing auxnet in the training for models trained with semi-supervision.

Manual examination of the segmentation results indicated that the main weakness of the weakly-supervised models is with the determination of the exact cell-cell boundary (supp. Fig. s1). Indeed we noticed that the weakly-supervised model can correctly infer the cell locations as well as the rough sizes and shapes of the cell, but has trouble discerning the detailed shape of the cell border. To correct this issue, we introduced an auxiliary convolutional network (auxnet) during the training specifically to predict the cell boundaries (Fig 2c). Of course the auxnet cannot be directly trained against ground truth, as the cell boundaries are not part of the annotation. Therefore, we train auxnet against the segmentation output from the main LACSS network (Fig. 2c), and the auxnet predictions in turn help to train the LACSS model weights by constraining LACSS outputs. Auxnet is only used during training and is not part of the computation during inference.

The concept of using one network to train another network can be found in many other self-supervised training techniques, e.g., generative adversarial network^30^ (GAN). Importantly, we purposely designed the auxnet to be of low field-of-view (FOV). The rationale is that this is also how humans perform cell segmentations. Typically we first scan a relatively large area in order to recognize the cell and understand its rough shape, but when it comes to tracing the exact cell boundaries, we will look at a much smaller area in order to find the exact pixel separating two cells. By designing the auxnet to be of low FOV, we force it to focus on a different set of image features than that of LACSS, which has a much larger FOV.

Introducing the auxnet in the training significantly improved the model accuracy (Fig. 4). For three of the cell lines (BV2, SKOV3 and SkBr3), the differences between the weakly-supervised models and fully-supervised models are within one percent point. For the rest of the cell lines, the weakly-supervised models still underperform, but the gaps are much smaller.

We also compare our results with the previously published baselines (Table 1) and find our results to be competitive. For BV2 and SkBr3 lines, our models trained with incomplete annotations actually had AP50 scores slightly higher than the previous models trained with the full annotations. When using more stringent IOU criteria (e.g. 75%) our results are slightly worse, which is expected because achieving high IOU often requires the model to have some knowledge of the annotation bias of the exact annotators, which our training pipeline has no access to. Our results showed the highest performance gap in the SHSY5Y cell line (Table 1). However, for this cell line, the LACSS model performs poorly even when trained under full supervision (AP_50_=0.520), suggesting that the weakness stems from the model architecture and not from the training strategy. One possibility is that for SHSY5Y, a neuroblastoma with particularly complex shapes, our model backbone (Resnet-50^31^) does not have sufficient expressibility compared to the baseline models, which are based on Resnet-200^31^.

**Table 1.**
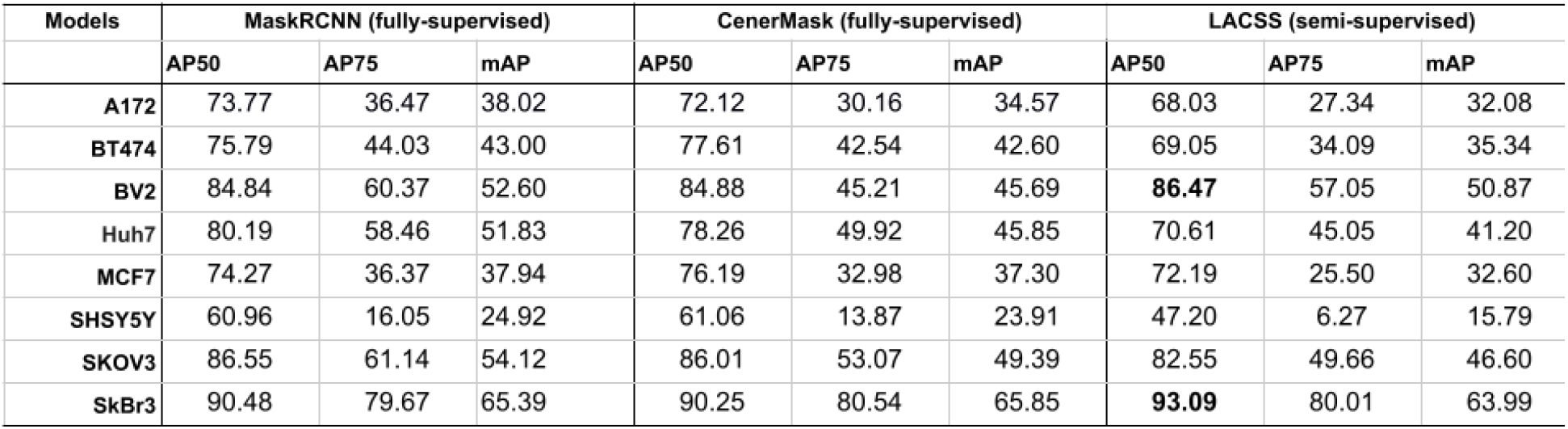
Comparison of LACSS model and the baselines. All baseline models were trained with the complete annotation, whereas LACSS models were trained using the incomplete annotation. Bold font indicated cases where the LACSS model performed better than the baselines.

**Table 2.** Benchmarking LACSS models trained with lab-generated image data and annotations. The training dataset includes 500 immunofluorescence images of A431 cells at high density. All annotations were derived from experimental data semi-programmably. See text for more details.

| IOU% | 50 | 55 | 60 | 65 | 70 | 75 | 80 | 85 | 90 | 95 | mAP/AFNR |
| --- | --- | --- | --- | --- | --- | --- | --- | --- | --- | --- | --- |
| AP | 0.8385 | 0.7981 | 0.7254 | 0.6267 | 0.4883 | 0.2813 | 0.0843 | 0.007 | 0 | 0 | 0.38496 |
| FNR | 0.1373 | 0.1665 | 0.2192 | 0.2878 | 0.3903 | 0.5532 | 0.7629 | 0.9325 | 0.9965 | 1 | 0.54462 |

Fig. 5 showed examples of segmentation results on the eight cell lines. It can be seen that the model generally was able to trace the boundary between cell-cell contacts, despite the fact the training data contains no such information. The model also to a degree recaptured certain annotation biases. For example, for BV2 cells, the annotator chose not to label dying cells with an abnormal appearance, and the model similarity avoided their detections. On the contrary, for BT474 cells, the annotator frequently avoided labeling cells at the center of the colonies, probably due to poor contrast. However, this preference was not learned by the model.

**Figure 5.**
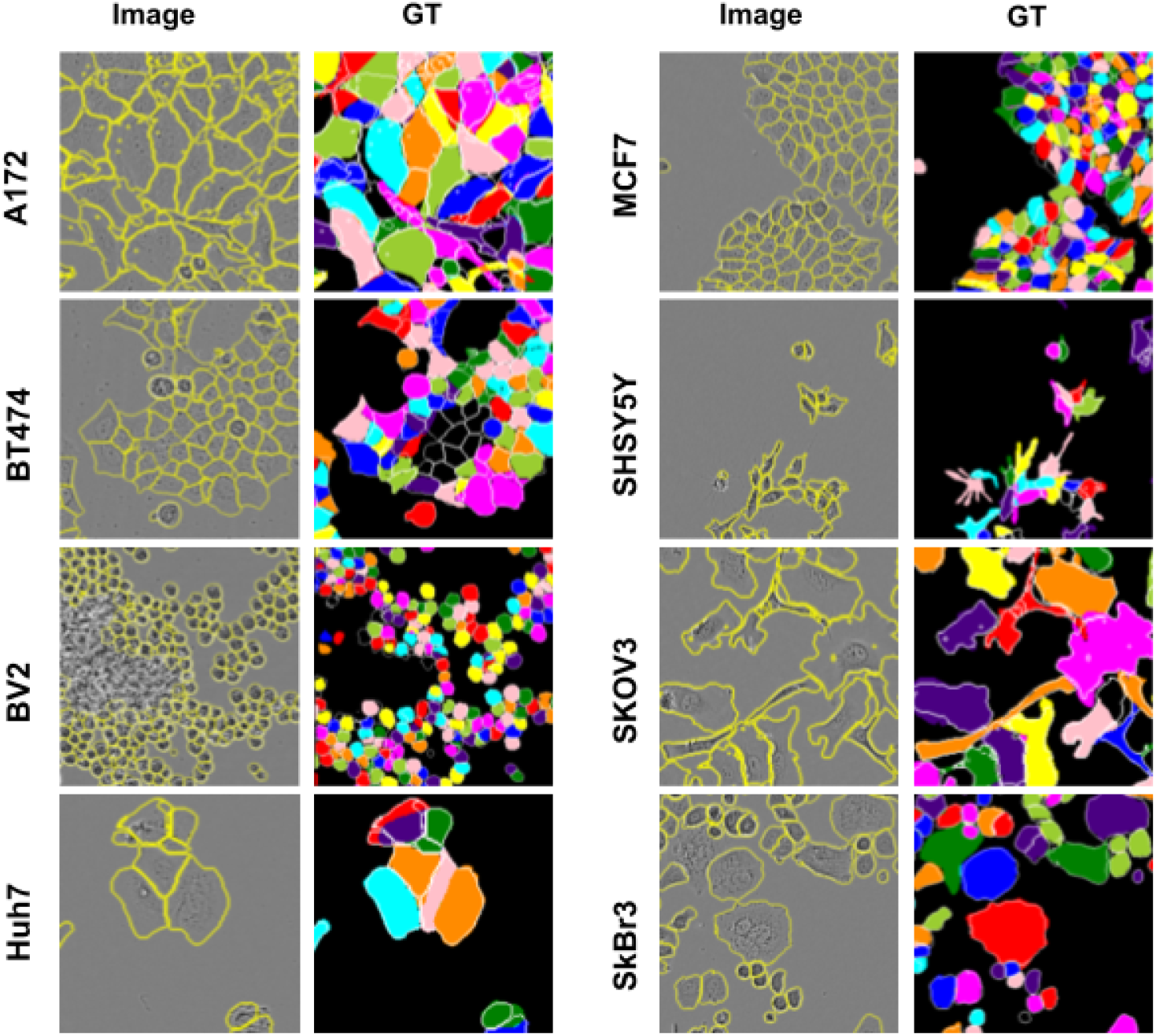
Segmentation examples for LIVECell dataset using LACSS models trained with the semi-supervision configuration. Examples for all eight cell lines were shown. The input phase contrast images (left) were overlaid with model predictions drawn as yellow contours. The ground truth segmentations were shown on the right, for which the model predictions were also shown as white contours for comparison.

### Efficient training pipeline with semi-automatic annotation

Finally we turn to test a training pipeline, in which the annotation is generated semi-programmably (Supp. Fig. s2). To do that, we acquired a dataset on cultured A431 cells using fluorescence microscopy. The cell line is chosen for its tendency to grow in large colonies with closely packed cells, which means that there will be significant loss of information going from the full segmentation annotation to the image-level segmentation. This allows for a more stringent test of our training strategy. The cells were immuno-fluorescently labeled using anti-PY100 antibody, which localize to both cell membrane and cell cytosol (Fig. 6), producing a useful contrast to identify individual cells. Additionally, nucleus images were acquired on a separate channel with DAPI staining. We acquired 500 images of 512×512 pixels using an automated microscope, which served as the training set. Furthermore, using the same protocol but in a separate experiment, we acquired 25 more fluorescence images, which we manually segmented to serve as the validation set.

**Figure 6.**
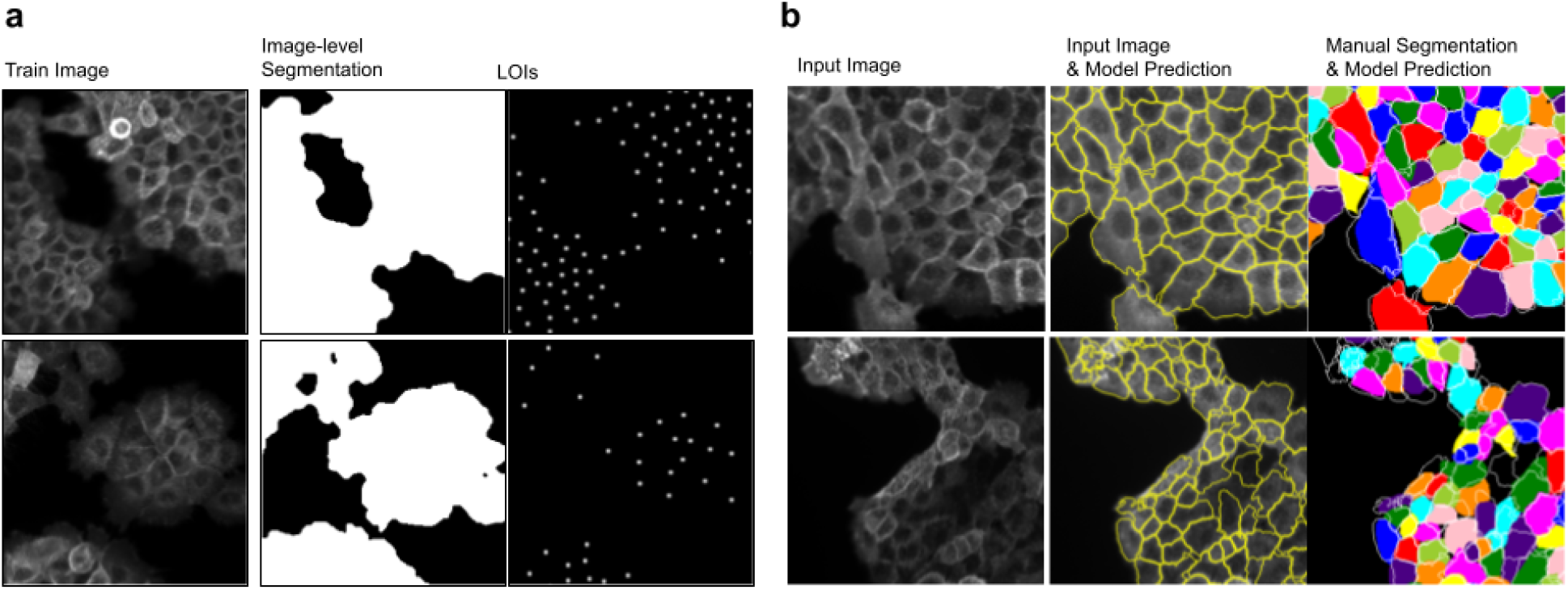
LACSS model trained with incomplete annotations. (**a**) Examples of training data and accompanying annotations generated semi-programmably. (**b**) Examples of model inferences showing input fluorescence image (left), the model prediction as yellow contours (middle), and comparison of model prediction with manual segmentation of the same input image (right). The bottom example represented a more challenging case with atypical image contrasts.

To generate the LOIs for the training set, we performed nucleus detection on the DAPI images. For this purpose, we used a very simple blob detection algorithm based on the difference-of-Gaussian filter, even though much more sophisticated algorithms are available^3,32^. This is to minimize the dependencies on external training data. Even with such a simple algorithm, the results were accurate enough: we estimated that both false positive and false negative rates to be around 1% based on the visual inspections. We also opted to *not* manually correct these mistakes, in order to evaluate the model performance in a more streamlined pipeline. To generate image-level segmentations, we used an existing graph-cut software to process all 500 fluorescence images. Because each image requires a slightly different threshold value for correct segmentation, we automatically generated five sets of segmentation results using a preset series of threshold values. An annotator then manually examined the results to pick one for each input image that is considered the best.

In Fig. 6 we showed representative examples of the segmentation results. The model shown here achieved an AP_50_ of 0.84 (Fig 6b), which we consider to be of practical use for various research tasks, such as single-cell analysis. The main advantage is that our model was established with a much faster speed than the traditional pipelines that relies on manually generated single-cell annotations. The result also confirms that the exact nature of the LOIs can be flexible. In previous tests, we used the locations of the center-of-mass of cell segmentations, because the dataset was already pre-segmented. Here the nucleus locations were typically off-center to the cell and randomly localized. Neither are the LOIs center of the nucleus, as the results depend on the exact intensity distributions of the DAPI signal. However, these features did not seem to prevent training of the segmentation model.

## Discussion

Annotation burden is a well-known problem of deep learning methods. The issue is particularly challenging for single-cell segmentation tasks. Most of the existing works dealing with this problem focused on the goal of reducing the amount of annotation as much as possible, which we believe may not be the most productive venue. Instead, we focus on utilizing annotations that can be generated automatically or semi-automatically, leveraging on experimental resources that are commonly available in biological research labs. This allows us to retain as much annotation information as possible, while not being slowed down by manual annotations. We believe this is the best chance to create an efficient model building pipeline, without sacrificing too much with regard to the model accuracy, which is one of the top priorities for most of the biological researchers.

In particular, we demonstrate here that segmentation models can be built by utilizing two types of weak annotations: the whole-image level segmentation and the location-of-interests, both of which are incomplete descriptions of the cell segmentation, but combined provided sufficient information to achieve good model accuracies. Indeed, we showed several examples, where models build our incomplete annotations performed on par with models trained with full supervision. On the other hand, even in cases where our method slightly underperforms, LACSS model may still be a good compromise due to the significantly reduced cost for the model building.

In conclusion, we provided a practical alternative for efficiently building single-cell segmentation deep-learning models, which would be of interest to biological researchers in need of performing single-cell analysis from microscopy images.

## Methods

### Image acquisition

For this study we experimentally acquired a small microscopy dataset on the human squamous-cell carcinoma line A431 (ATCC CRL1555). Cells were seeded on glass substrate and grown to ~80% confluence in standard growth media before fixation for imaging. Cells were labeled with the standard immunofluorescence protocol: They were washed with a blocking buffer (2% BSA in PBS, 1% TX100), incubated with Alexa647-labeled anti-PY100 (1:500 in blocking buffer containing DAPI) for 2 h while rocking at 4°C. Images were acquired on an automated inverted fluorescence microscope (Olympus IX) with an 20x objective. Images were captured on a camera with a 512×512 sensor. The per pixel dimension is 0.8 micrometer under this configuration.

### Image annotation

Only the lab-acquired A431 cell data were annotated by us. All other dataset used in this study was previously segmented manually. To annotate the A431 training set, we use the existing GraphCut function of the FIJI software to produce image-level segmentation. All images were segmented at five different intensity thresholds and a human annotator later manually examined the output to pick one out of five for each input image. To produce LOIs, we first apply a difference-of-Gaussian filter to the DAPI images, with the σ values of 4.2 and 5 pixels. We then searched for all local maxima with four-connectivity and used their locations as LOIs. To annotate images in the validation set, we used the online annotation software described by Stringer et. al^2^ to generate single-cell segmentations for all cells in the image. The software takes polygon input indicated by mouse clicks and converts the inputs into single-cell segmentations.

### Model training

Here we provide a general outline of the model training procedure. See suppl. text for detailed descriptions of the model architectures, the hyperparameter choices and the training procedures of each model.

Models for the cell image library dataset were all trained with random initialization of model weights. The experiments were repeated five times and the best model was chosen based on the highest AP_50_ score on the validation set. Models for LIVECell dataset were also trained with random initialization of weights, i.e., without pretraining on external datasets. Best models were chosen based on benchmarks on the validation set. The testing set was not involved in any manner for choosing models. For the lab-acquired dataset, the model was pretrained on the LIVECell dataset, then continuously trained on the lab-acquired A431 cell training data and evaluated on the validation set. All models were trained using a single NVIDIA Tesla A100 GPU. ADAM optimizer was used for all experiments.

### Model benchmarks

To evaluate model accuracy, we relied primarily on the average precision (AP) metrics, which were widely used in the instance segmentation literature.

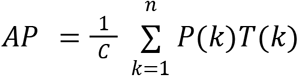

where *C* is the total number of ground truth cells, *n* is the total number of detections, *T*(*k*) is a indicator function of whether the *k*-th detection is positive (*T*(*k*) = 1) or negative (*T*(*k*) = 0), and *P*(*k*) is the precision of the first *k* detections. Whether a detection is considered positive is determined by the intersection-over-union (IOU) of the detection against the ground truth. We use the notation AP_IOU_ to denote the AP values at a specific IOU threshold, i.e, AP_50_ is the average precision when the positive detection requires a minimal IOU of 50%. We also use the notation mAP to denote the average AP over a series of IOU thresholds. In our study, we chose IOU thresholds from 0.5 to 0.95 (inclusive) in a 0.05 step size. Per convention of instance detection literature, we do not allow multiple detections to match against the same ground truth instance.

A secondary metric we used is the false negative ratio (FNR), which is simply

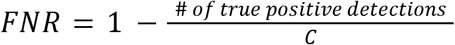

The metric measures the percent of cells that were missed by the model. As in AP metric, we used the notation *FNR_IOU_* to distinguish *FNRs* at different IOU thresholds, and we used *AFNR* to denote average FNR at all thresholds.

## Code availability

All software source code is available at https://github.com/jiyuuchc/lacss

## Notes

https://github.com/jiyuuchc/lacss

### Competing Interest Statement

The authors have declared no competing interest.

