## supplemental information for "Efficient End-to-end Learning for Cell Segmentation with Machine Generated Incomplete Annotations"

### Supplementary text

#### LACSS model architecture

LACSS model contains three sub-modules, all of which are based on fully-convolutional neural network (FCN) designs.

Backbone. The model takes an image as input, represented by a tensor of (H×W×C), where H and W are the size of the image and C is the number of color channels. The backbone module of the model extracts feature representations of the image at multiple resolutions. In this study, we employed a ResNet50 based backbone. The encoder of the backbone is a modified ResNet50 with added channel attention mechanism. The original ResNet50 employs a stem component that performs a downsampling operation, which we eliminated in our model. Instead, our stem is simply two layers of convolution filters with ReLU activation, resulting in a tensor of (H×W×64). We do this because the ResNets were originally designed and optimized for the ImageNet dataset. The typical objects in ImageNet are larger than a typical cell in our microscopy images. Other researchers resolve the scale mismatch by pre-scaling the input image. Our solution is slightly more memory efficient. In addition, we modified the ResNet50 by adding channel attention operations at the end of each major processing block. Channel attention is a light-weight mechanism that has been shown to increase model expressibility with little added computation cost.

Our decoder is a standard feature pyramid network[1] (FPN). For a fixed input size, the ResNet encoder outputs representations at 2x, 4x, 8x and 16x down-sized dimensions, corresponding to output level 1-4. The FPN integrates these features via convolution and addition to produce new integrated outputs at the same four levels. We used the same channel number (256) for outputs at all levels. We call the output with the largest dimension (level 1) high-resolution feature, which will serve as the input to the segmentation network. We call the rest low-resolution features, which feed into the location proposal network (LPN) (see Fig. 1).

LPN. The LPN is analogous to the regional proposal network[2] (RPN) of the popular MaskRCNN model for instance segmentation. Its task is to predict a set of LOIs from the input image. It follows a simple design of a multi-layer convolutional network. For a input feature tensor with the dimension H/s $\times$ W/s, the LPN outputs a tensor (**S**) of dimension (H/s, W/s, 1), representing scores of H $\times$ W / $s^2$ potential location proposals. Here s is the downscaling factor of the features, that is either 4, 8, or 16. The elements of the score tensor corresponding to a 2D grid of locations in the original image separated by s pixels. To determine the exact positions of the proposal LOIs, the LPN also outputs a regression tensor (**R**) of the shape (H/s, W/s, 2), representing the offset from the grid locations. We used the sigmoid activation for the score tensor and used no activation for the regression output.

While the LPN is very similar to RPN, there are two important differences. Firstly, Unlike RPN, which computes regression for the object bounding-box (location and size), the LPN does not predict the object size, because the information is unavailable during weakly-supervised training. Secondly, RPN typically takes features at all scales as inputs. This allows RPN to separate larger object detection from small object detection. The larger objects were localized using features of small-dimensions and smaller objects were from features of large-dimensions. In our system, however, we do not have information regarding object size. Therefore, LPN computes on only one scale of features (i.e, level 2, 3 or 4), pre-selected as a hyperparameter of the model. This indeed limits the ability of LACSS to detect objects of different sizes. However, we believe the impact is relatively minor, because unlike generic object detection tasks where object sizes were influenced by factors such as camera perspective, most cell segmentation problems deal with a narrower size distribution.

Combining the score and regression output, we can produce a ranked list of location proposals. We perform a simple clustering of the proposal locations based on the non-max-suppression algorithm to remove redundant proposals that are very close together. After that, for each proposed location $(ii,\ jj)$ we generate a location tensor (***L***) based on a simple location encoding:

$$L_{ijk} = \begin{cases} i - ii & \text{if } k = 0 \\ j - jj & \text{if } k = 1 \end{cases}$$

We have also experimented with more sophisticated location encoding schemes, including using learned encodings, and found them to be not helpful with regard to the model performances.

An extra step was taken during training (but not during inference) to mix the predicted locations with the ground truth locations. The mix operation starts with the list of ground truth locations, and replacing any ground truth location with a proposal location if the two are sufficiently close (i.e., below a preset threshold distance). The resulting mixture locations were used to generate the location tensors instead of the predicted LOIs.

Segmentation FCN. The segmentations for individual cells were computed from the level-one features. The feature tensor was first concatenated with the location tensors produced by the LPN, resulting in multiple copies of hybrid tensors, each will result in one segmentation prediction. The segmentation network is a simple multi-layer fully-convolutional network (FCN). To increase the computational efficiency and reduce memory consumption, the first few convolution layers ignore the location tensor part of the data, therefore the computational results can be shared for all location inputs. The location tensor is only used in computation at the last two convolutional layers. In addition, before feeding the data to the last two layers, we crop the input tensor to a smaller patch surrounding the respective LOIs, and batch all data together to increase the parallelism of the computation. This is simply because in almost all cases, cells are of limited sizes and there is no need to compute for every pixel of the image. The size of the crop is a preconfigured hyperparameter of the model. Because the input feature for the segmentation network is down-scaled from microscopy image by 2x, the last layer of the segmentation network uses a transposed convolution operation, which rescales the segmentation results back to original resolution. We used ReLU activation for all layers, except the last one, which used the sigmoid activation.

Auxnet. Auxnet is an additional component that is only utilized during semi-supervised training, and not in fully-supervised training or in inference. It takes the microscopy image as the input and attempts to predict all cell boundaries. It is designed simply as a four-layer convolutional network with no down-sampling or pooling operations, so that the overall field-of-view (FOV) of the network is kept low.

### Fully-supervised end-to-end learning

LACSS is capable of fully-supervised end-to-end learning using the standard stochastic gradient descent (SGD) optimization. The model loss function in this configuration has two components: the LPN loss and the segmentation loss.

LPN loss. As described earlier, the LPN outputs two tensors representing the scores (**S**) and offset regressions (**R**) with regard to a 2D grid of locations in the original image. To construct the LPN loss function, we first compute two additional tensors representing the ground truth scores ($\mathbf{S_T}$) and ground truth offsets ($\mathbf{R_T}$), using the provided ground truth annotations. The $\mathbf{S_T}$ was set to be 1 if the corresponding grid location is within 1.5**s** distance away from a ground truth LOI; otherwise the $\mathbf{S_T}$ was set to be 0. To construct $\mathbf{R_T}$, we first find for each grid location the nearest LOI among all ground truth LOIs. We then compute the offsets of each grid location to its nearest LOI, which becomes the $\mathbf{R_T}$.

We then used focal cross-entropy loss[3] to measure the difference between **S** and $\mathbf{S_T}$, and we used Huber loss to measure the difference between **R** and $\mathbf{R_T}$. The LPN loss is then the sum of the two. In addition, to avoid noisy offset predictions at grid locations far away from any LOIs, we masked off all elements where $\mathbf{S_T}$ was 0 when computing the Huber loss. More formally,

$$L_s = -\alpha(1-p_t)^\gamma \log p_t;\ p_t = \begin{cases} S & \text{if } S_T = 1 \\ 1-S & \text{if } S_T = 0 \end{cases}$$

$$L_R = \begin{cases} \frac{1}{2}(R-R_T)^2 & \text{if } (R-R_T) < \delta \\ \delta \cdot (|R-R_T| - \frac{1}{2}\delta) & \text{otherwise} \end{cases}$$

and

$$L_{LPN} = \langle L_S \rangle + \langle L_R \rangle$$

The bracket denotes average over all positions. We used the fixed values of $\alpha = 0.25$, $\gamma = 2.0$ and $\delta = 1.0$ for all models in this study.

Segmentation loss. We use cross-entropy loss between the predicted segmentations (**G**) and ground truth segmentations ($\mathbf{G_T}$) to measure segmentation loss:

$$L_{FS-SEG} = \langle -G_T \log G - (1 - G_T)\log(1 - G) \rangle$$

#### Weakly-supervised end-to-end learning

Under the semi-supervised configuration, the LACSS model loss has three components: the LPN loss, the weakly-supervised segmentation loss and the cell boundary loss.

LPN loss. The computation of the LPN loss is identical to that of the fully-supervised configuration, because the ground truth LOIs were provided as part of the annotations.

Weakly-supervised segmentation loss. Since we no longer have the instance level ground truth segmentation (**$G_T$**), we have to construct a differentiable loss function using the incomplete annotations:

$$L_{WS-SEG} = \left\langle -C \cdot G^{[i]} - (1 - C) \cdot (1 - G^{[i]}) - G^{[i]} \sum_{k \neq i} \log(1 - G^{[k]}) \right\rangle_i$$

Here we use a superscript on $G^{[i]}$ to denote the i-th instance prediction from the model and the bracket average is over both the locations and the instances.

The first half of the loss function checks for the consistency between the single-cell segmentations with the image-level segmentation (**C**). The formulation,

$$- C \cdot G^{\{i]} - (1 - C)(1 - G^{[i]})$$

was somewhat parallel to the more commonly used cross-entropy loss function, which would have been of the form

$$- C \log G^{[i]} - (1 - C)\log(1 - G^{[i]}).$$

Both formulations are minimized at the same point of G=C. Additionally, they have the same gradient at this minimal point. However, by replacing the log term with a linear term, our formulation has a much shallower gradient at locations where G≠C. The intention is to allow the segmentation output to be different from **C**, by not incurring exceedingly high losses.

The second half of the loss function checks the consistency between individual segmentation instances. The term $-G^{[i]}\sum_{k\neq i}\log(1-G^{[k]})$ is minimized when either $G^{[i]}=0$ or $G^{[k]}|_{k\neq i}\equiv 0$. In other words, this part of the loss function aims to minimize the overlaps between individual instances. We have previously shown that a loss function of this type shown can be used to separate a cluster of cells into individual instances in a self-supervised manner. Here we employ this technique again for the end-to-end training of the segmentation model.

Cell boundary loss. The LACSS model performs significantly better when we train it with an additional loss term that specifically focuses on accurate cell boundary calculation. Since cell boundaries are not available from the annotation, we predict them using an auxiliary network (auxnet) directly from the input image, yielding a prediction $\mathbf{B}_P$. At the same time, we also computed the cell boundary locations from LACSS output using a bit of heuristics,

$$\mathbf{B}_H = \tanh\left[\sum_i \phi\left(\mathbf{G}^{[i]}\right)\right],$$

where $\phi()$ is the sobel edge filter and the hyperbolic tangent function (tanh) is applied to ensure the final results were bound between (0, 1), similar to $\mathbf{B}_P$. The cell boundary loss is the Huber loss between $\mathbf{B}_H$ and $\mathbf{B}_P$:

$$L_{cb} = \begin{cases} \frac{1}{2}(B_P - B_H)^2 & \text{if } (B_P - B_H) < 0.5 \\ \frac{1}{2}\left|B_P - B_H\right| - \frac{1}{8} & \text{otherwise} \end{cases}$$

We train the auxnet by minimizing the cell boundary loss $L_{cb}$.

We train LACSS model by minimizing the sum of all losses:

$$L_{LACSS} = w_0 L_{LPN} + w_1 L_{WS_SEG} + w_2 Lcs$$

where $w_i$ are loss weights. in this study we used only constant weights ($w_i \equiv 1$) for all models.

#### LACSS Experiments

Cell Image Library. All input images were normalized by scaling the gray values to the range of [0, 1]. Total of 89 images were used for training and the rest for validation. We used level-3 features as inputs for LPN. The maximum crop size for segmentation is 128 pixels, corresponding to a 64x64 patch size for the level-1 feature inputs of the segmentation FCN. We augment the training dataset by random flipping (both horizontal and vertical) and random resizing (10% both up and down).

To train the LACSS models with full supervision, we used the original manual segmentation label and trained the model continuously for 10,000 steps. The validation metrics are frequently monitored (every 250 steps) to closely monitor the overfitting issues. The model weights were randomly initialized and training was repeated five times to pick the best model.

To train the LACSS models with weak-supervision, we converted the original single-cell segmentation label to image-level segmentations and LOIs (center of mass). Model weights were randomly initialized and we ran the training for 20,000 steps, recording validation metrics every 1,000 steps. Training was repeated five times. We did not include auxnet during training for this dataset.

LIVECell. Dataset was downloaded from the publisher's website. All image data were normalized by scaling the gray values to the range of [0, 1]. This dataset was imaged with a slightly lower magnification, therefore we used a lower segmentation crop size of 96 pixels. We were also concerned that Resnet50 may not be well-performing on detecting very small cells. Therefore we resized images of the two smallest cell lines, BV2 and MCF7, by 2 foldes. In addition, a subset of cells from the SKOV3 line are exceptionally large and exceed the 96p crop size. Therefore we scaled all images of SKOV3 down by 30%. We employed the same data augmentation as in Cell Image Library, i.e, using random flipping and random resizing. All train images were then either cropped or zero-padded to a fixed size (544 x 704) before input to the model.

We initialized model weights randomly in both the supervised and the semi-supervised configurations. We did some preliminary comparisons study by initializing the ResNet component of the model with weights pretrained on ImageNet and found the results to be not significantly different. Therefore, all results reported in this study were based on random weights.

For both supervised and semi-supervised training, we trained the models for 105000 training steps, checking $AP_{50}$ of the validation set every 3500 steps and picked the model based on the best $AP_{50}$. We then perform the full benchmarking on the testing set with the picked model. One hyperparameter of the model is the feature scale used for LPN. We trained models on all three choices (level 2, 3 and 4) and compared the results. For semi-supervised training, we found that the best model was cell-line dependent. For example, SHSY5Y cells achieved a much better benchmark when using level-2 features and MCF7 cells were much better with level-3. Therefore we chose different model hyperparameters for different cell lines. On the other hand, for fully-supervised training, we found no significant differences when altering this parameter. Therefore we only reported results from the model using level-3 features for LPN.

Lab-generated immuno-fluoresence dataset. Only semi-supervised models were generated for the lab dataset, since we did not fully segment the training images. The models used level-3 features for LPN and a segmentation crop size of 128 pixels. Unlike the previous cases, the training images intensity were rescaled normally, with a mean of 0 and variance of 1.0, because the fluorescence images had a much heavier tail in the intensity distributions. We again employed random flipping and random resizing for data augmentation.

We used the transfer learning strategy for this dataset. The model was pre-trained on the LIVECell dataset with full-supervision. We then train the model on the fluorescence dataset for additional 30,000 steps with semi-supervision. The pretraining allows for much faster convergence, and therefore less training steps were needed on the fluorescence data.

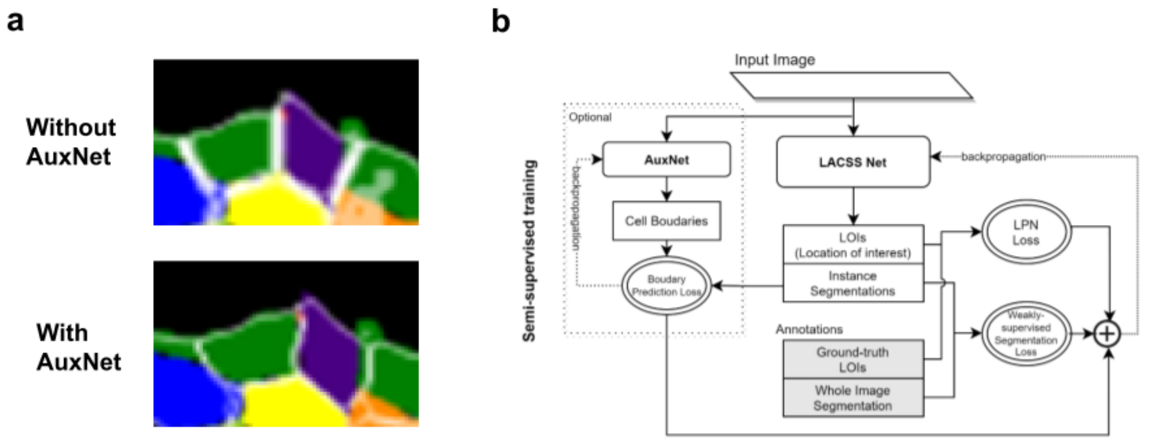


**Figure S1**. Comparison of segmentation results with and without auxnet. **(a)** Without auxnet (top), the LACSS model has defects in tracing the cell boundary in pixel-level accuracy. Adding auxnet (bottom) in the training pipeline (bottom) significantly improved the accuracy of cell boundaries. **(b)** Diagram outlining training algorithm with and without auxnet.

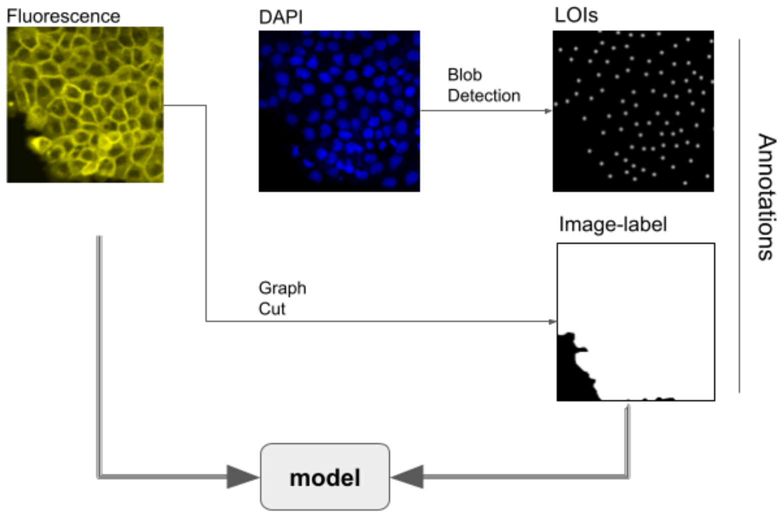


**Figure S2.** Schematic showing the streamlined annotation procedure using experimental data.
